# promor: a comprehensive R package for label-free proteomics data analysis and predictive modeling

**DOI:** 10.1101/2022.08.17.503867

**Authors:** Chathurani Ranathunge, Sagar S. Patel, Lubna Pinky, Vanessa L. Correll, Shimin Chen, O. John Semmes, Robert K. Armstrong, C. Donald Combs, Julius O. Nyalwidhe

## Abstract

**Summary:** We present promor, a comprehensive, user-friendly R package that streamlines label-free (LFQ) proteomics data analysis and building machine learning-based predictive models with top protein candidates.

**Availability and implementation:** promor is freely available as an open source R package on the Comprehensive R Archive Network (CRAN)(https://CRAN.R-project.org/package=promor) and distributed under the Lesser General Public License (version 2.1 or later). Development version of promor is maintained on GitHub (https://github.com/caranathunge/promor) and additional documentation and tutorials are provided on the package website (https://caranathunge.github.io/promor/).

## 1 Introduction

Label-free quantification (LFQ) approaches are fast becoming popular in mass spectrometry-based proteomics. One of the most widely used software tools for protein identification and quantification is MaxQuant (Tyanova *et al*., 2016*a*). The downstream analysis of MaxQuant output files can be complex and often challenging to those inexperienced in proteomics data analysis. Some tools available for this purpose are implemented as graphical user interface (GUI) applications, among which, one of the most popular is the MaxQuant-associated tool, Perseus (Tyanova *et al*., 2016*b*). Perseus is an extensive software suite that offers a range of features to analyze several different types of of proteomics data. While Perseus is fairly easy to use, the user interface with its wide range of options can be overwhelming at times to new users. Further, the inability to save previously used analytical settings in Perseus leaves more room for human error and makes it more difficult and time consuming to reproduce previous results and figures. Other tools such as MSstats (Choi *et al*., 2014), are primarily implemented as R packages, and provide greater analytical flexibility and reproducibility to proteomics data analysis workflows. While these available software all offer analytical capability to perform the steps in typical proteomics data analysis workflows, users may need additional software to perform tasks specific to their research domain (e.g. clinical applications, biomarker discovery).

In recent years, machine learning (ML) has made its presence felt in the field of proteomics. Particularly in biomarker research, ML has become a necessary last step to derive candidate biomarker panels from proteomics data. ML algorithms are now being widely employed to build proteomics-based predictive models of disease prognosis and diagnosis. When building a proteomics-based predictive model, choosing a robust panel of protein candidates can greatly improve the accuracy of the model. In this regard, ML-based predictive models could benefit from narrowing down protein features to those that show significant differences in abundance between groups of interest. In the current landscape of proteomics data analytical tools the capability to seamlessly transition from differential expression analysis to predictive modeling is lacking. Realizing this need, we developed promor, a comprehensive, user-friendly, R package that streamlines differential expression analysis and predictive modeling of label-free proteomics data. promor provides an all-in-one reproducible workflow that integrates tools to perform quality control, visualization, and differential expression analysis of label-free proteomics data. Further, promor provides tools to build ML-based predictive models using top protein candidates identified through differential expression analysis, assess model performance, determine feature importance, and estimate the predictive power of the models.

## 2 Overview

### 2.1 Implementation

promor is implemented in R (≥ 3.5.0) and relies on packages such as imputeLCMD (Lazar *et al*., 2016), limma (Ritchie *et al*., 2015), and caret (Kuhn, 2008) for back-end pre-processing, differential expression analysis, and machine learning-based modeling, respectively. As input, promor requires a user-generated tab-delimited text file containing the experimental design and a MaxQuant-produced “proteinGroups.txt” file or a standard quantitative table of protein intensities, which could be produced by any proteomic data anlaysis software. For visualization, promor employs the popular ggplot2 (Wickham *et al*., 2016) architecture and produces ggplot objects, which allows for further customization (Figure 1).

**Fig. 1.**
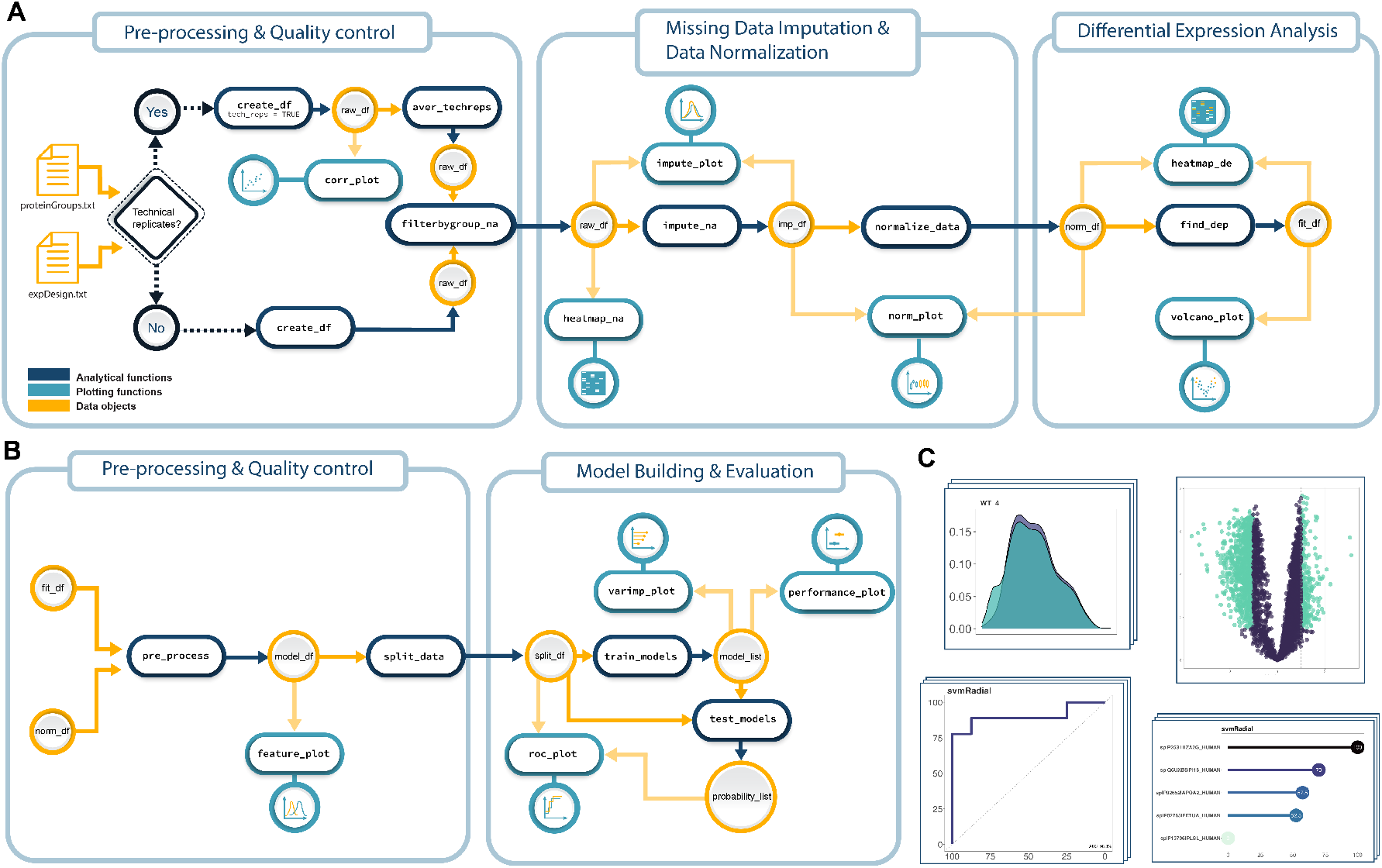
An overview of suggested promor workflows. (A) Proteomics data analysis workflow includes analytical functions for pre-processing, quality control, missing data imputation, data normalization, and differential expression analysis. (B) Modeling workflow includes analytical functions for pre-processing the output of differential expression analysis, model building, and model evaluation. (C) Several plotting functions are provided to visualize data and produce publication-ready figures using color blind-friendly palettes.

### 2.2 Proteomics data analysis

promor can be used to analyze any bottom-up label-free proteomics data (e.g. raw, LFQ, or iBAQ). Multiple functions are provided for quality control, visualization, missing data imputation, normalization, and differential expression analysis (Figure 1A), (Table 1).

**Table 1.** Analytical and visualization functions in promor

| Function name | Input | Tasks | Output |
| --- | --- | --- | --- |
| create_df | A proteinGroups.txt file from MaxQuant / a standard input file containing a quantitative matrix of protein intensity data.<br>A tab-delimited text file containing the experimental design. | Creates a data frame of LFQ protein intensities. Removes contaminant proteins, proteins identified only by site, reverse sequence proteins, and proteins identified by less than two unique peptides. Converts zeros to missing values. Log<sub>2</sub> transforms the values. | raw_df |
| aver_techreps | raw_df | If technical replicates are present in the data, computes average intensity across technical replicates for each sample. | raw_df |
| filterbygroup_na | raw_df | Filters out proteins with more than 34% missing values (less than 66% valid values) in at least one of the groups. | raw_df |
| impute_na | raw_df / norm_df | Imputes missing values using the "minProb" method. | imp_df |
| normalize_data | raw_df / imp_df | Normalizes the data using the "quantile" method. | norm_df |
| find_dep | norm_df / imp_df | Identifies differentially expressed proteins with an absolute log<sub>2</sub> fold change > 1 at an adjusted p-value < 0.05. | fit_df |
| pre_process | norm_df / imp_df<br>fit_df | Extracts protein intensity data for the top 20 differentially expressed proteins, removes proteins that show high pairwise correlation (> 0.90), and converts the data into a format suitable for modeling. | model_df |
| split_data | model_df | Splits data into training (70%) and test (20%) data-sets while preserving the overall class distribution of the data. | split_df |
| train_models | split_df | Trains ML models on the training data-set using the default list of ML algorithms ("svmRadial", "glm", "rf", "xgbLinear"), performs 10-fold cross validation three times, calculates re-sampling-based performance measures for the models, and outputs the best model for each algorithm. | model_list |
| test_models | split_df<br>model_list | Uses the models built using the training data to predict the test data | probability_list |
| corr_plot | raw_df | Generates scatter plots showing the correlation between pairs of technical replicates. | ggplot |
| heatmap_na | raw_df | Generates a heatmap to show the missing data distribution in the matrix. | ggplot |
| impute_plot | raw_df / norm_df<br>imp_df | Generates a global density plot showing the data distribution before and after missing data imputation. | ggplot |
| norm_plot | raw_df / imp_df<br>norm_df | Generates box plots showing the sample data distributions before and after data normalization. | ggplot |
| heatmap_de | imp_df / norm_df<br>fit_df | Generates a heatmap of protein intensities for the top 20 differentially expressed proteins. | ggplot |
| volcano_plot | fit_df | Generates a volcano plot highlighting significantly differentially expressed proteins (absolute log<sub>2</sub> fold change > 1 at an adjusted p-value < 0.05). | ggplot |
| feature_plot | model_df | Generates box plots showing protein intensity differences among groups (classes). | ggplot |
| varimp_plot | model_list | Generates lollipop plots showing the importance of different proteins (features) in the models built. | ggplot |
| performance_plot | model_list | Generates boxplots showing the performance (accuracy and kappa) of models built using different ML algorithms. | ggplot |
| roc_plot | split_df<br>probability_list | Generates Receiver Operating Characteristic (ROC) curves showing the predictive power of the models built using different ML algorithms. | ggplot |

\*This table describes the tasks and the output produced by the functions under default settings.

To demonstrate the utility of promor for analyzing label-free proteomics data that do not contain technical replicates, we analyzed a previously published proteome benchmark data-set by Cox *et al*. (2014) (PRIDE ID: PXD000279). The data-set consists of LFQ protein intensity data for 6694 proteins quantified from HeLa (H) and E. coli (L) lysates that were mixed at defined ratios. There were six samples in total. Three biological replicates represented each of the two groups. The results from the analysis were visualized at multiple stages (Supplementary Figures S1 - S5). First we pre-processed the data using the `create_df` function with default settings. `create_df` function removed contaminant proteins, proteins identified ‘only-by-site’, reverse sequence proteins, and proteins identified by fewer than two unique peptides. To remove proteins with a high proportion of missing values, we used the `filterbygroup_na` function, setting the highest allowed missing data percentage in either group at 40%. Next, we imputed the missing data in the data frame using the `impute_na` function with the default ‘minProb’ method assuming that the missing values are left-censored. Since the data have already been normalized with the MaxLFQ algorithm Cox *et al*. (2014) in MaxQuant we did not further normalize the data in promor. The output of imputation (`imp_df` object) was used in the differential expression analysis, performed using the default settings in the `find_dep` function. We identified 1294 siginificantly differentially expressed proteins between the ‘H’ and ‘L’ groups in the data (Table S1, Figures S4 and S5).

Further, to test the utility of promor for analyzing label-free proteomics data that contain technical replicates, we analyzed previously published data by Ramond *et al*. (2015)(PRIDE ID: PXD001584). This data-set consists of LFQ protein intensity data obtained from two strains (WT - wild type and D8 - ΔargP mutant) of *Francisella tularensis*, a pathogenic bacterium responsible for the zoonotic disease tularemia. The proteinGroups.txt file contained LFQ data for 1265 proteins across 18 samples representing the two conditions (WT and D8) with three biological replicates in each condition, and three technical replicates for each biological replicate. A step-by-step tutorial providing a detailed description of the workflow and the implementation choices are provided here: https://caranathunge.github.io/promor/articles/promor_with_techreps.html

### 2.3 Building predictive models

In promor, multiple functions are provided to build predictive models with differentially expressed proteins and assess model performance (Figure 1B),(Table 1). Over 200 machine learning algorithms are made accessible through the caret package (Kuhn, 2008) for building predictive models. For users inexperienced in complex machine learning algorithms, promor provides a default list of five widely used classification-based algorithms, chosen to represent a variety of machine learning model types (e.g. random forest, support vector machines, generalized linear models, naive bayes, and gradient boosting). However, while many different algorithms can be applied to proteomics data, it is important to note that not all of them are well-suited to address the problem at hand. The choice of machine algorithms should be carefully decided according to the prediction task, data type, sample size, and the number of features (proteins) in the data-set.

We tested the use of promor for building predictive models by analyzing a previously published data-set by Suvarna *et al*. (2021) (PRIDE ID: PXD022296). In the original study the authors built proteomics-based classification models to predict COVID severity in patients. To avoid class imbalance in the data, only a subset of the samples were used from the original proteinGroups.txt file. The steps leading up to differential expression analysis are described in detail here: https://caranathunge.github.io/promor/articles/promor_for_modeling.html. The results from differential expression analysis (`fit_df` object) and the normalized data frame (`norm_df` object) were used in the modeling workflow. The `fit_df` and `norm_df` objects were pre-processed with the `pre_process` function to convert the data into a `model_df` object. Next, we split the data into training and test data-sets using the `split_data` function. The training data-set contained 70% of the data (29 samples), while the test data-set contained the remaining 30% (6 samples). The `train_models` function was run on the training data-set in the `split_df` object with four selected ML algorithms: Random Forest (rf), Support Vector Machine with Linear Kernel (svmLinear), Naive Bayes (naive_bayes), K-Nearest Neighbor(knn). The four algorithms were chosen based on their suitability for building models using few features (eight proteins) and samples (35 samples). Further, a k-fold cross validation (k = 10, repeats = 3) was employed to evaluate model performance. The output was used to test the models on the test data-set included in the `split_df` object. The results from the analysis were visualized at multiple levels during the modeling workflow (Figures S6 - S9). The model built with the ‘naive_bayes’ algorithm performed best in terms of accuracy (85.5) and AUC (88.9%)(Figure S9).

### 2.4 Benchmarking

We compared the performance of promor against Perseus using the previously mentioned Cox *et al*. (2014) (PRIDE ID: PXD000279) data-set. An identical workflow and parameters to those mentioned in the section 2.2 were used in Perseus. In Perseus, we used the imputeLCMD plugin to implement the “minProb” imputation method, and the limma plugin to implement the moderated t-test. We observed a significant overlap in the differentially expressed proteins identified by both programs (98.85%) (Figure 2A). The number of proteins that were only identified by a single program could be attributed to the random sampling during missing value imputation. Further, the calculated log-fold changes and p-values were strongly correlated between the two programs (Figure 2B,C). R code for benchmarking analysis is provided on github at https://github.com/caranathunge/promor_bioRxiv_preprint

**Fig. 2.**
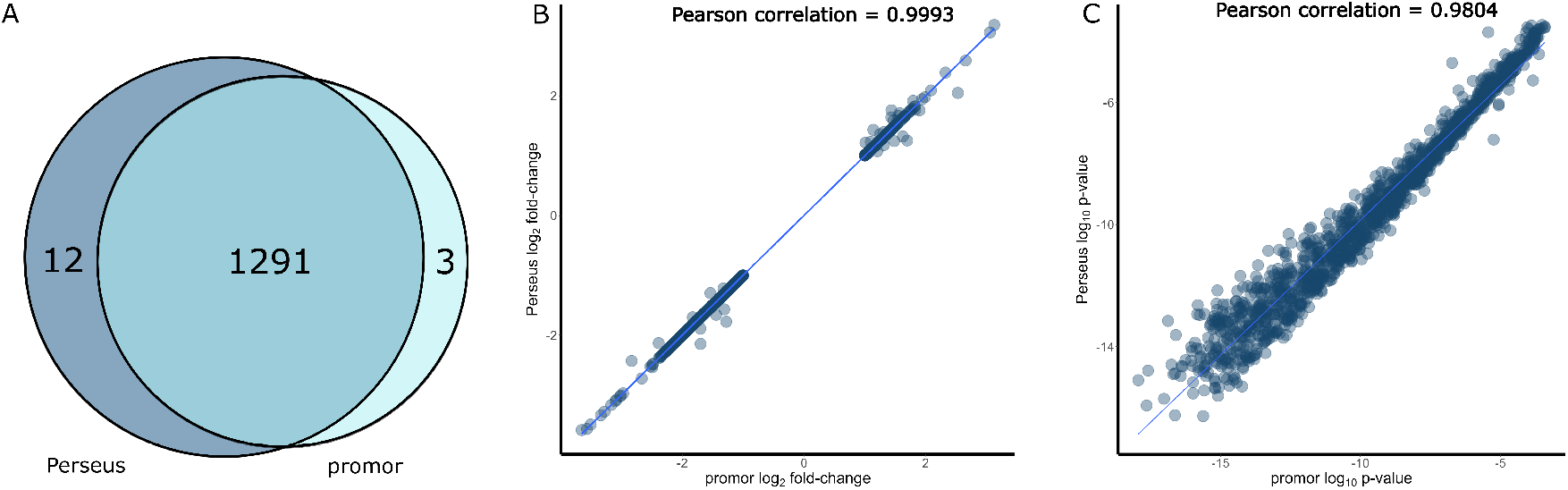
A comparison between promor and Perseus using the proteome benchmark data-set, Cox et al. (2014). (A) A Venn diagram showing the overlap of the significantly differentially expressed proteins identified by promor and Perseus. Scatterplots of the resulting protein log_2_ fold changes (B) and log_10_ p-values (C) of differentially expressed proteins as calculated by promor and Perseus.

## 3 Conclusions

We present promor, a user-friendly, comprehensive R package that facilitates seamless transition from differential expression analysis of label-free proteomics data to building predictive models with top protein candidates; a feature that could be particularly useful in clinical and biomarker research.

## 4 Acknowledgements

We wish to thank Asitha I. Senanayake for his helpful comments and discussions on software development.

## References

Choi, M., Chang, C.Y., Clough, T., Broudy, D., Killeen, T., MacLean, B. and Vitek, O. (2014) Msstats: an r package for statistical analysis of quantitative mass spectrometry-based proteomic experiments. Bioinformatics, 30 (17), 2524–2526.

Cox, J., Hein, M.Y., Luber, C.A., Paron, I., Nagaraj, N. and Mann, M. (2014) Accurate proteome-wide label-free quantification by delayed normalization and maximal peptide ratio extraction, termed maxlfq. Molecular & cellular proteomics, 13 (9), 2513–2526.

Kuhn, M. (2008) Building predictive models in r using the caret package. Journal of statistical software, 28, 1–26.

Lazar, C., Gatto, L., Ferro, M., Bruley, C. and Burger, T. (2016) Accounting for the multiple natures of missing values in label-free quantitative proteomics data sets to compare imputation strategies. Journal of proteome research, 15 (4), 1116–1125.

Ramond, E., Gesbert, G., Guerrera, I.C., Chhuon, C., Dupuis, M., Rigard, M., Henry, T., Barel, M. and Charbit, A. (2015) Importance of host cell arginine uptake in francisella phagosomal escape and ribosomal protein amounts*[s]. Molecular & Cellular Proteomics, 14 (4), 870–881.

Ritchie, M.E., Phipson, B., Wu, D., Hu, Y., Law, C.W., Shi, W. and Smyth, G.K. (2015) limma powers differential expression analyses for rna-sequencing and microarray studies. Nucleic acids research, 43 (7), e47–e47.

Suvarna, K., Biswas, D., Pai, M.G.J., Acharjee, A., Bankar, R., Palanivel, V., Salkar, A., Verma, A., Mukherjee, A., Choudhury, M. et al. (2021) Proteomics and machine learning approaches reveal a set of prognostic markers for covid-19 severity with drug repurposing potential. Frontiers in physiology,, 432.

Tyanova, S., Temu, T. and Cox, J. (2016a) The maxquant computational platform for mass spectrometry-based shotgun proteomics. Nature protocols, 11 (12), 2301–2319.

Tyanova, S., Temu, T., Sinitcyn, P., Carlson, A., Hein, M.Y., Geiger, T., Mann, M. and Cox, J. (2016b) The perseus computational platform for comprehensive analysis of (prote) omics data. Nature methods, 13 (9), 731–740.

Wickham, H., Chang, W. and Wickham, M.H. (2016) Package ‘ggplot2’. Create elegant data visualisations using the grammar of graphics. Version, 2 (1), 1–189.

