## Supplemental Material for "promor: a comprehensive R package for label-free proteomics data analysis and predictive modeling"

Supplementary Material

Chathurani Ranathunge, Sagar S. Patel, Lubna Pinky, Vanessa L. Correll, Shimin Chen, O. John Semmes, Robert K. Armstrong, C. Donald Combs, and Julius O. Nyalwidhe

#### 1 Supplementary Figures

Note: Code used for producing the following figures is available on GitHub (`https://github.com/caranathunge/promor_bioRxiv_preprint/blob/main/promor_supplementary_figures.Rmd`)

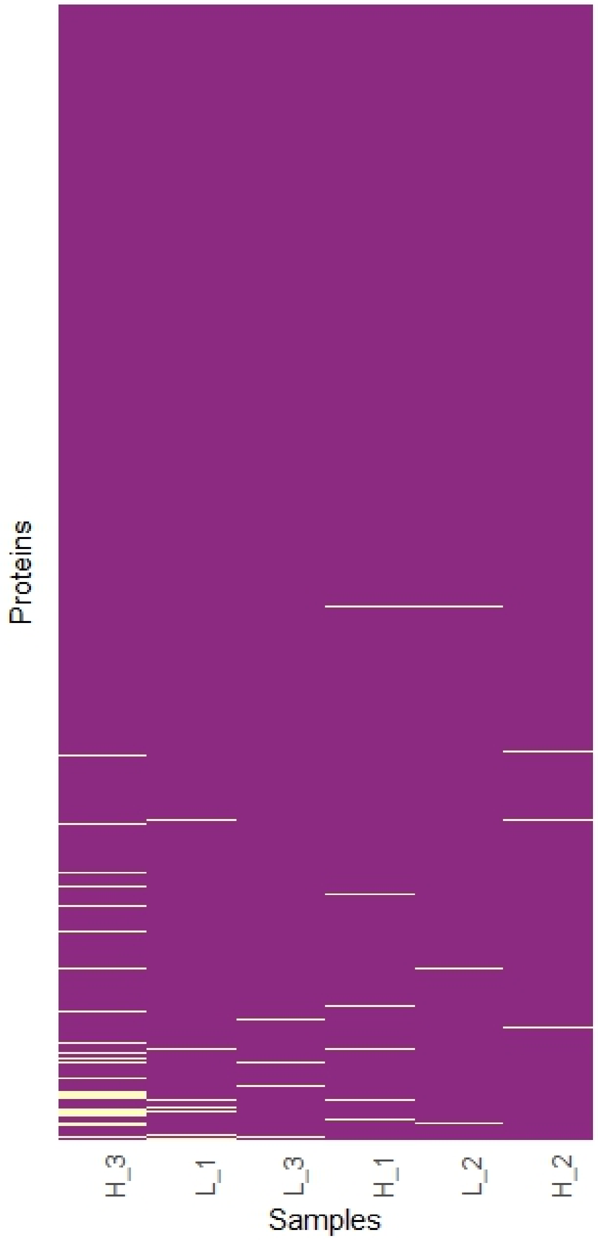


Fig. S 1: A heatmap showing the missing data distribution in protein intensity data. The data consists of protein intensity values for 4584 protein groups (rows) in six samples. The proteins on the y-axis have been reordered by mean intensity and the samples on the x-axis by the sum of intensity to show missing data distribution patterns in the data. Proteins with lower mean intensity appear to have higher levels of missing data and sample H_3 has the most number of missing values. Input data from Cox *et al.* (2014) downloaded from the PRIDE Archive (PRIDE ID: PXD000279).

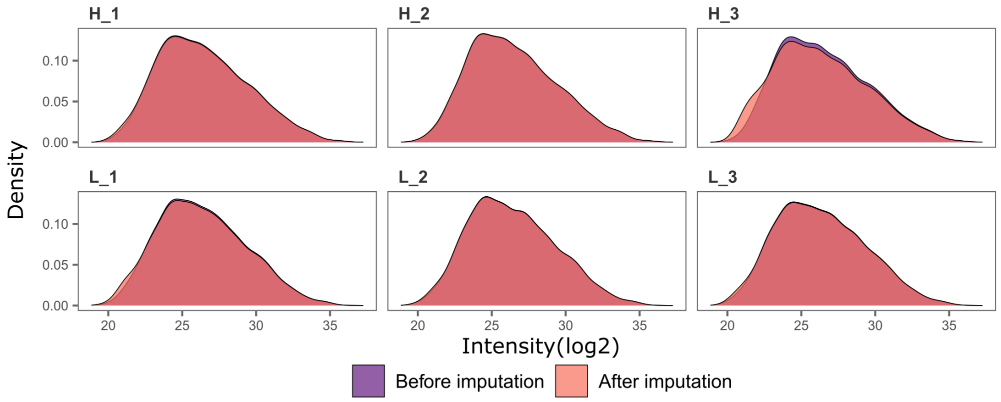


Fig. S 2: Density plots showing the impact of missing data imputation on the protein intensity data distribution in each sample. ‘minProb’ imputation method was used. Input data from Cox *et al.* (2014) downloaded from the PRIDE Archive (PRIDE ID: PXD000279).

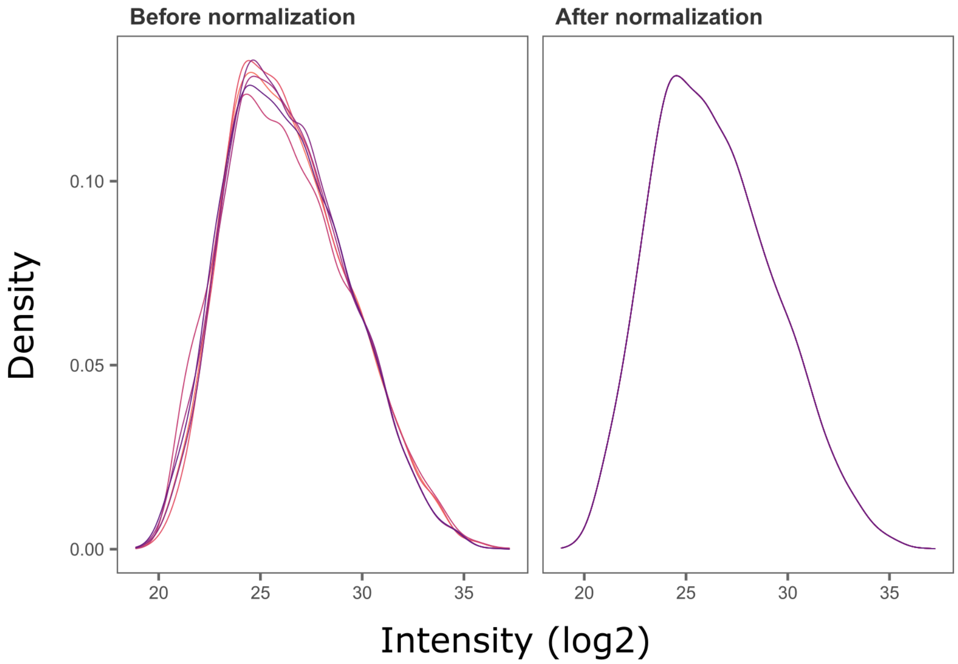


Fig. S 3: Density plots showing the impact of data normalization on the protein intensity data distribution. ‘quantile’ normalization method was applied here for visualization purposes only. Input data from Cox *et al.* (2014) downloaded from the PRIDE Archive (PRIDE ID: PXD000279).

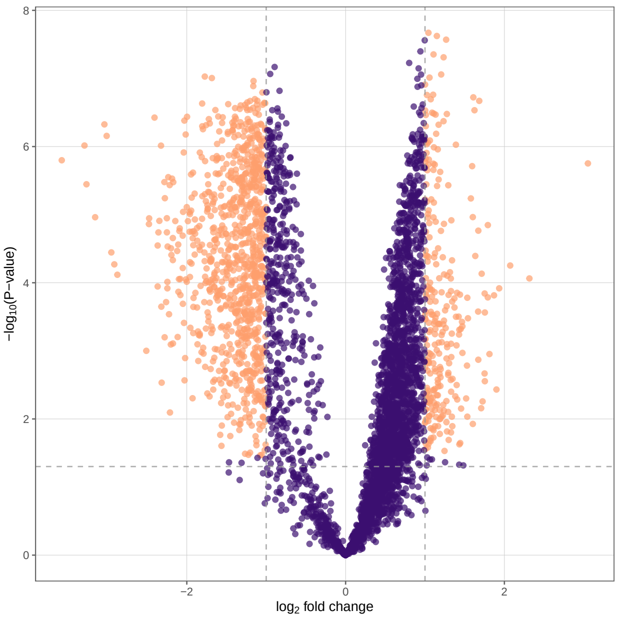


Fig. S 4: Volcano plot showing the results from the differential expression analysis. Significance criteria used: log-2-fold change of 1 and adjusted p-value of 0.05. Protein groups meeting the significance criteria are colored in orange. Input data from Cox *et al.* (2014) downloaded from the PRIDE Archive (PRIDE ID: PXD000279).

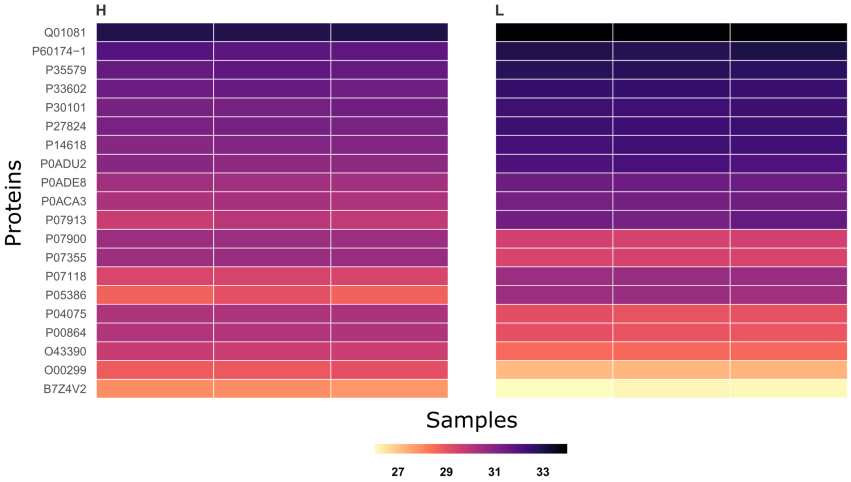


Fig. S 5: Heatmap showing the results from the differential expression analysis. The top 20 protein groups that meet the significance criteria (log-2-fold change of 1 and adjusted p-value of 0.05) are shown here. Input data from Cox *et al.* (2014) downloaded from the PRIDE Archive (PRIDE ID: PXD000279).

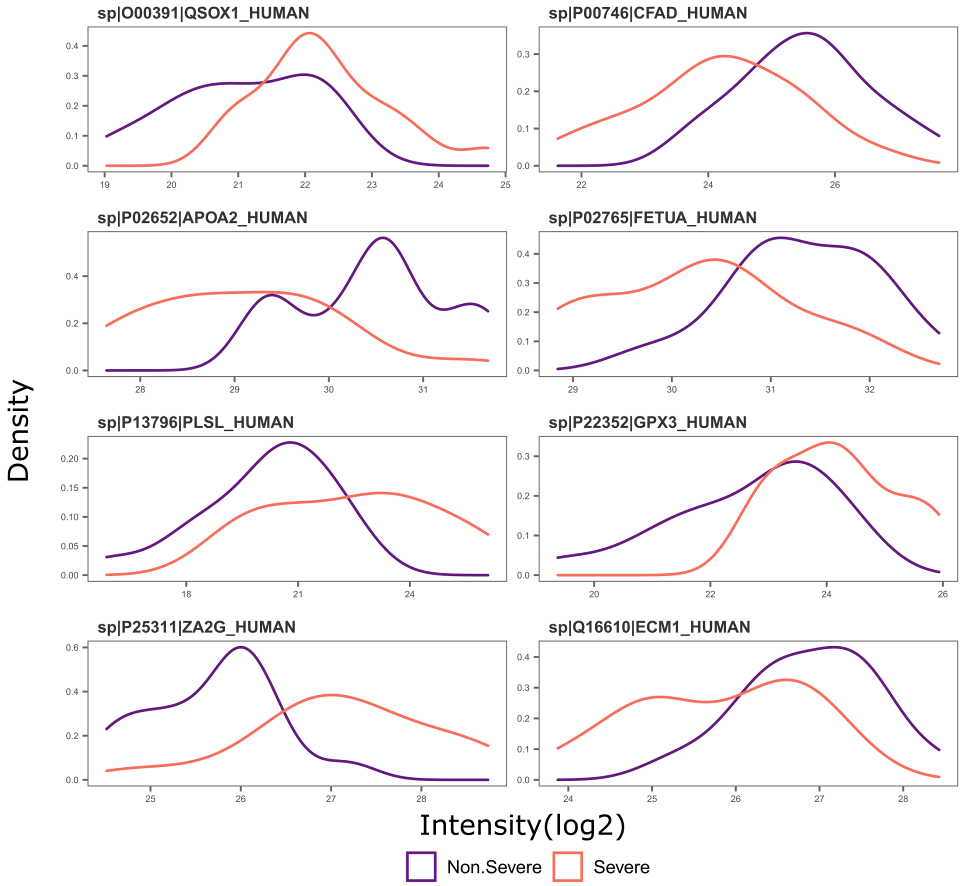


Fig. S 6: The variation of intensities in six proteins between COVID severe and non-severe patient groups. Input data from Suvarna *et al.* (2021) downloaded from the PRIDE Archive (PRIDE ID: PXD022296).

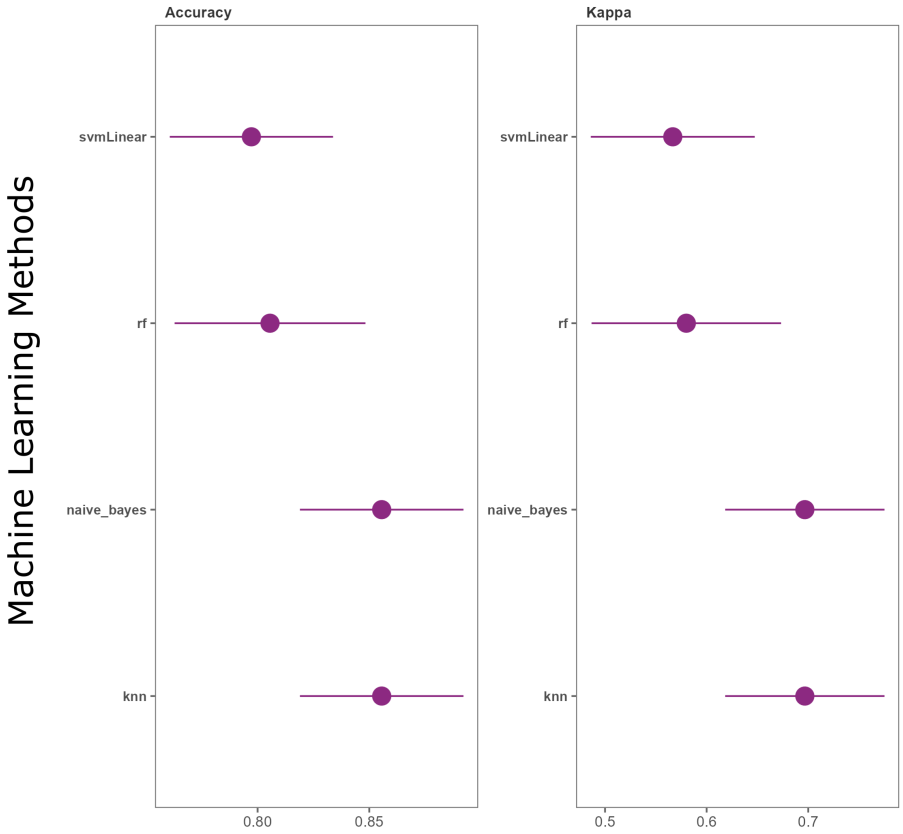


Fig. S 7: Dot plots showing model performance. Models built using four machine-learning algorithms are assessed for performance using Accuracy and Kappa. Input data from Suvarna *et al.* (2021) downloaded from the PRIDE Archive (PRIDE ID: PXD022296).

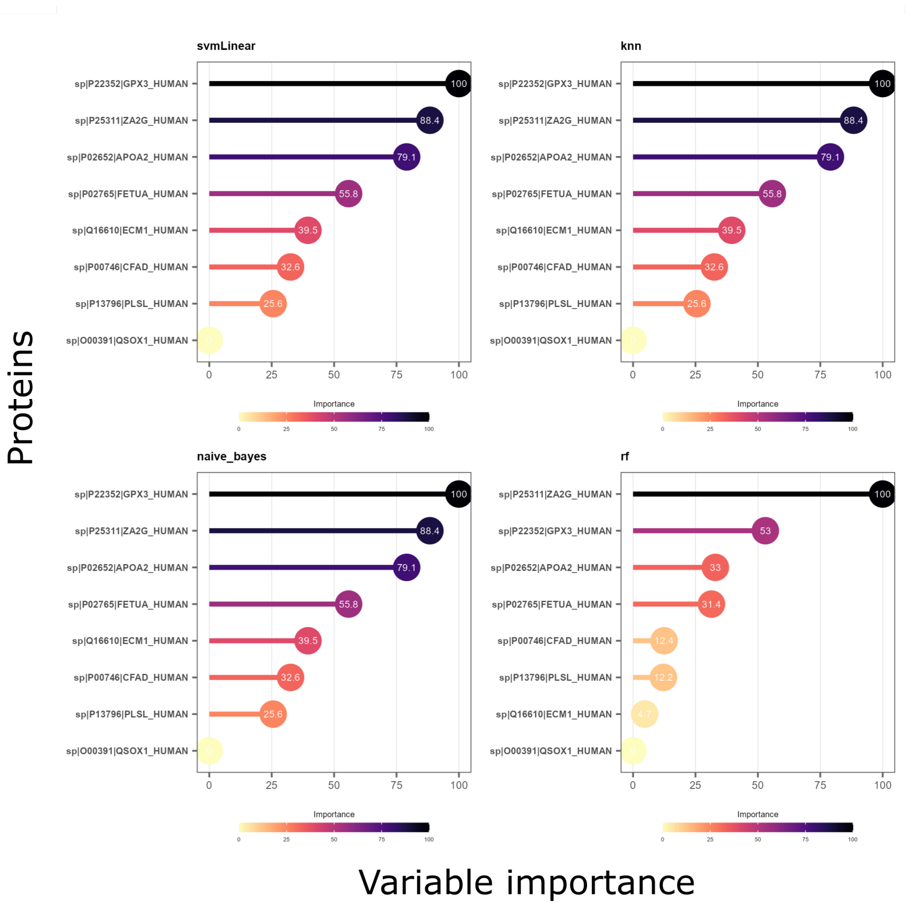


Fig. S 8: Lollipop plots showing variable (protein) importance. Proteins are sorted according to their importance in the models built using four different machine learning algorithms. Input data from Suvarna *et al.* (2021) downloaded from the PRIDE Archive (PRIDE ID: PXD022296).

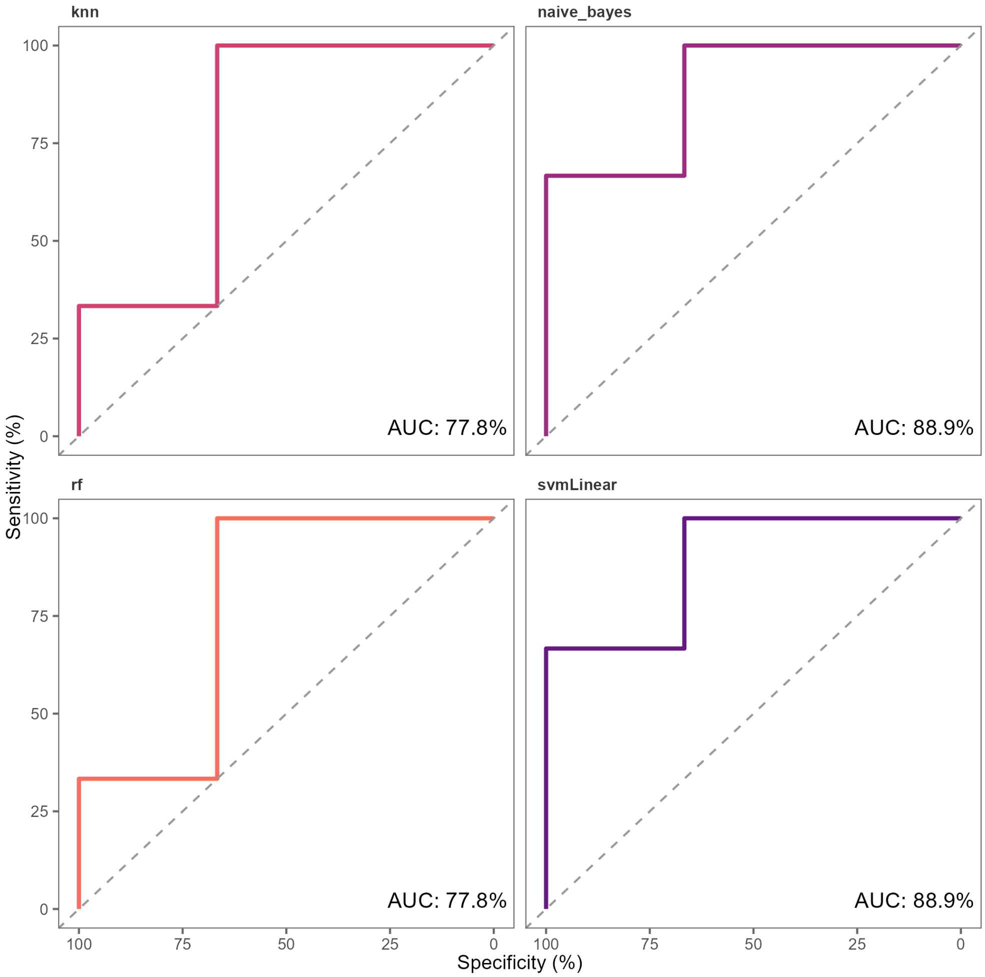


Fig. S 9: Receiver operating characteristic curves illustrating the diagnostic ability of the models built using four different machine learning algorithms. Area under the curve (AUC) estimates are indicated in each plot. Input data from Suvarna *et al.* (2021) downloaded from the PRIDE Archive (PRIDE ID: PXD022296).

### References

Cox,J., Hein,M.Y., Luber,C.A., Paron,I., Nagaraj,N. and Mann,M. (2014) Accurate proteome-wide label-free quantification by delayed normalization and maximal peptide ratio extraction, termed maxlfq. *Molecular & cellular proteomics,* **13** (9), 2513–2526.

Suvarna,K., Biswas,D., Pai,M.G.J., Acharjee,A., Bankar,R., Palanivel,V., Salkar,A., Verma,A., Mukherjee,A., Choudhury,M. *et al.* (2021) Proteomics and machine learning approaches reveal a set of prognostic markers for covid-19 severity with drug repurposing potential. *Frontiers in physiology,* **, 432.**
